# Structural insights into AT-rich DNA recognition by SALL family proteins

**DOI:** 10.1101/2022.06.13.496009

**Authors:** Wenwen Ru, Tomoyuki Koga, Xiaoyang Wang, Qiong Guo, Micha Gearhart, Shidong Zhao, Mark Murphy, Hiroko Kawakami, Dylan Corcoran, Jiahai Zhang, Zhongliang Zhu, Xuebiao Yao, Yasu Kawakami, Chao Xu

## Abstract

Spalt-like 4 (SALL4) plays an essential role in controlling the pluripotent property of embryonic stem cells (ESCs) via binding to AT-rich regions of genomic DNA. Here we present crystal structures of the the zinc finger cluster 4 (ZFC4) domain of SALL4 (SALL4^ZFC4^) bound with different double stranded DNAs containing a conserved AT-rich motif. In the structures, two zinc fingers of SALL4^ZFC4^ coordinatively recognize an AATA tetranucleotide. We also solved the DNA-bound structures of SALL3^ZFC4^ and SALL4^ZFC1^. These structures illuminate a common recognition mode for AT-rich DNA by the SALL family proteins. The DNA binding activity is essential for SALL4 function as DNA-binding defective mutants of mouse *Sall4* failed to repress aberrant gene expression in *Sall4*^-/-^ mESCs. Thus, these analyses provide new insights into the mechanisms of action underlying SALL family in controlling cell fate via preferentially targeted to AT-rich sites within genomic DNAs during cell differentiation.

## Introduction

Transcription factors (TFs) play essential roles in embryo development through binding to the specific regions of genomic DNA to direct different complexes in mediating programmable gene transcription ^1-3^. The occupancy of sequence-specific TFs is typically determined by the base composition within a genomic DNA region ^4-6^. It has been well known that unmodified CpG dinucleotide serves as signaling motif to recruit epigenetic regulators containing CXXC domain, a known CpG-binding module^7-9^. Although AT-rich regions are also highly enriched in some important regulatory genomic DNA elements, including TATA box^10, 11^, whether they also recruit sequence-specific TFs and how they function in embryo development are largely unknown ^12, 13^. Very recently, two lines of work independently identified Spalt-like transcription factor 4 (SALL4) as the AT-rich DNA binding protein via pull-down mass spectrometry screen and protein binding microarray, respectively^14, 15^. SALL4 belongs to the Spalt-like transcription factors (SALLs) family, which includes SALL1-4 ^16^. SALL4 is highly expressed in embryonic stem cells (ESCs) and several tumors, but absent in most adult tissues. Dysfunctional SALL4 pathway is associated with severe human diseases, including Holt-Oram syndrome (HOS)^17^, acro-renal-ocular syndrome (AROS)^18^, leukemogenesis and other cancers ^19^. SALL4 contains eight zinc fingers and they form four clusters ^14^. The zinc finger cluster 4 (ZFC4) is essential for SALL4 to recognize AT-rich sequence to repress expression of a variety of genes, and its mutation results in abnormal differentiation and embryonic lethality ^14, 15^. All SALL proteins except SALL2 recognize AT-rich DNAs via ZFC4 ^14, 15^. Despite the critical role of SALL4 ZFC4 in embryo development and its conservation in other SALL proteins, how it binds to AT-rich DNAs and how the SALL4 occupancy at AT-rich regions influences gene expression, remain elusive.

By using isothermal titration calorimetry (ITC) binding assay we ascertained that the ZFC4 of SALL3 and SALL4 prefer AT-rich DNAs. Then we solved the structures of SALL3 and SALL4 ZFC4 bound with different AT-rich DNAs, and revealed that the conserved ZFC4 specifically recognizes AT-rich motif via hydrogen bonding and van der waals interactions. In addition, we found that SALL4 ZFC1 also serves as a binder of AT-rich DNAs, albeit with weaker binding affinity. Inspired by previous finding that loss of Sall4 in ESCs causes aberrant neural gene expression ^20^, we evaluated the functional relevance of DNA-binding and found that DNA-binding deficient SALL4 mutant fails to repress aberrant expression of several genes, such as Irx3 and Irx5. Therefore, our study not only unveils how ZFC4 of SALL proteins preferentially recognizes AT-rich DNAs, but also sheds light on the biological importance of their binding to AT-rich DNA sequences in ESCs.

## Results

### ZFC4 of SALL4 and SALL3 selectively recognize AT-rich DNAs

To quantitatively study the binding activity of SALL4 to DNAs with different base compositions, we cloned, expressed, and purified the human SALL4^ZFC4^, spanning 856-930 (**Fig. 1a**), and measured its binding affinities towards different double stranded DNAs (dsDNAs) using ITC. Consistent with previous studies ^14, 15^, SALL4^ZFC4^ binds to different 12-mer DNAs containing AATATT with KDs in a range of 6.9-9.0 μM (**Fig. 1b-c, Supplementary Table S1**). In contrast, the binding was abolished when central four (ATAT) or six (AATATT) nucleotides were replaced by CG-rich nucleotides (**Supplementary Table S1**). To understand whether other SALL family members possess similar DNA binding selectivity, we cloned, expressed, and purified the ZFC4 domain of SALL3 (SALL3^ZFC4^) spanning aa 1102-1167, and examined its DNA binding property by ITC. Our binding analyses indicate that SALL3^ZFC4^ binds to the AATATT-containing 12-mer DNA with a KD of 8.0 μM (**Fig. 1d**), comparable to that of SALL4^ZFC4^ (KD = 6.9 μM). Like SALL4^ZFC4^, SALL3^ZFC4^ displayed no binding affinity towards CGCG-or CGCGCG-containing DNAs (**Supplementary Table S1**). Thus, we conclude that ZFC4 domains of SALL3 and SALL4 specifically recognize AT-rich DNAs judged by an *in vitro* binding assay.

**Figure 1.**
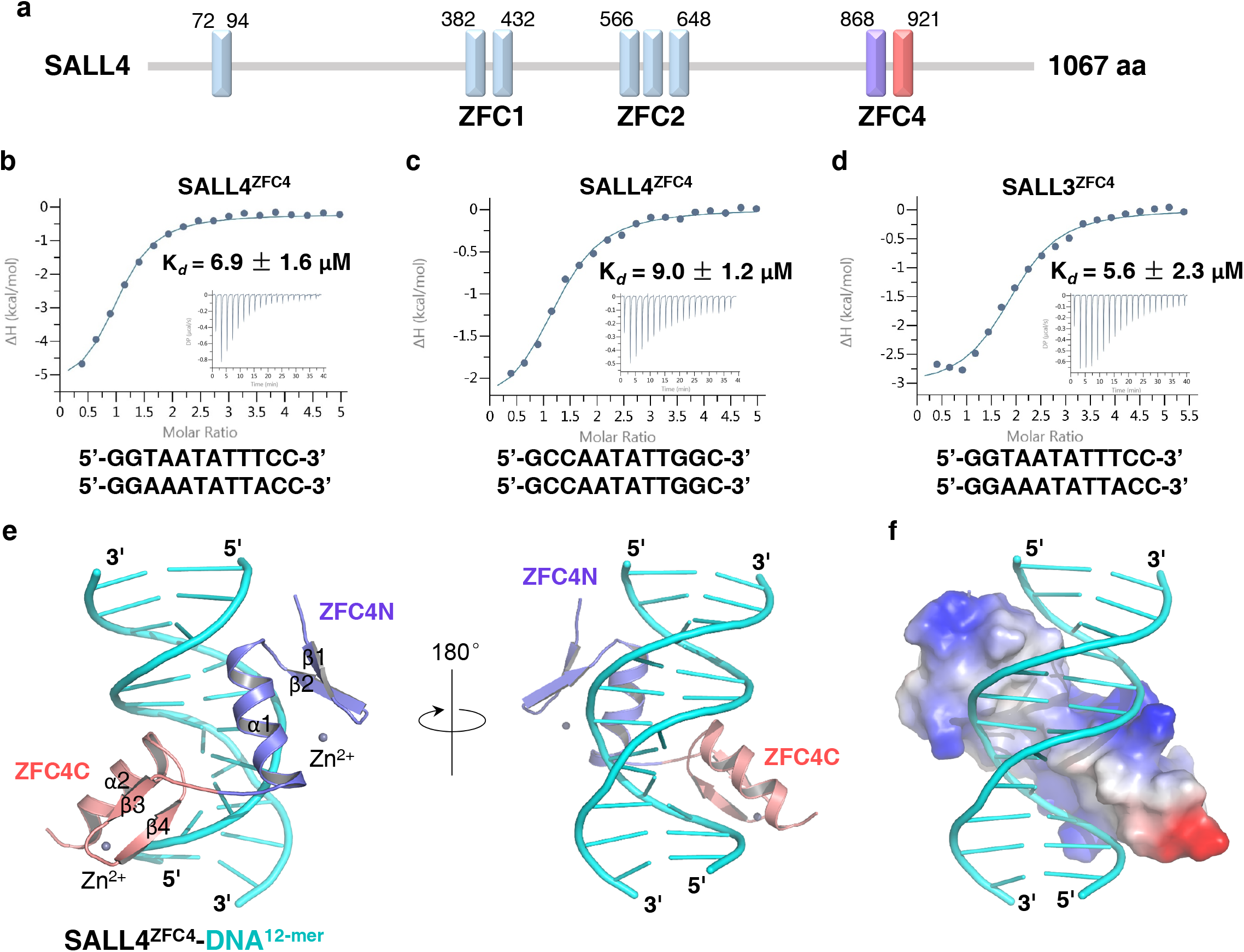
Structure of SALL4^ZFC4^ bound with the 12-mer dsDNA. **a** Domain architecture of human SALL4 containing ZFC1, ZFC2, and ZFC4. **b** ITC binding curves for SALL4^ZFC4^ binding to the 12-mer dsDNA (5’-GGTAATATTTCC-3’). **c** ITC binding curves for SALL4^ZFC4^ binding to the 12-mer dsDNA (5’-GCCAATATTGGC-3’). **d** ITC binding curves for SALL3^ZFC4^ binding to the 12-mer dsDNA (5’-GGTAATATTTCC-3’). **e** Crystal structure of SALL4^ZFC4^ with the 12-mer dsDNA (5’-GGTAATATTTCC-3’). The DNA is shown in cyan cartoon. Two zinc fingers of SALL4^ZFC4^, ZFC4N and ZFC4C, are shown in purple and salmon cartoon, respectively with indicated secondary structures. **f** Electrostatic surface of SALL4^ZFC4^ bound to the 12-mer dsDNA (cyan cartoon).

### The structures of SALL4 with different AT-rich DNAs

To gain insight into the molecular mechanism underlying AT-rich DNA recognition by SALL4^ZFC4^, we solved the crystal structure of the SALL4^ZFC4^ with a 12-mer dsDNA (5’-GGTAATATTTCC-3’) in a 2.45 Å resolution (**Supplementary Table S2**). There are two double-stranded DNA molecules in a asymmetric unit, with each of them bound with two SALL4^ZFC4^ molecules. SALL4^ZFC4^ is comprised of two zinc fingers, termed as ZFC4N and ZFC4C **(Fig. 1e, Fig. 2a**). In each complex, one SALL4^ZFC4^ molecule binds to the central major groove of the dsDNA with both zinc fingers visible, whereas the other one binds to the end of the dsDNA with only ZFC4C visible. Given that ITC binding data suggest that SALL4^ZFC4^ binds to the 12-mer dsDNA in a molecular ratio of 1:1 (**Supplementary Table S1**), binding of the second SALL4^ZFC4^ to the dsDNA is likely due to the crystal packing, and the invisible ZFC4N might be due to its intrinsic flexibility. Therefore, our structural analysis focuses on the SALL4^ZFC4^ molecule bound at the central major groove of the dsDNA.

**Figure 2.**
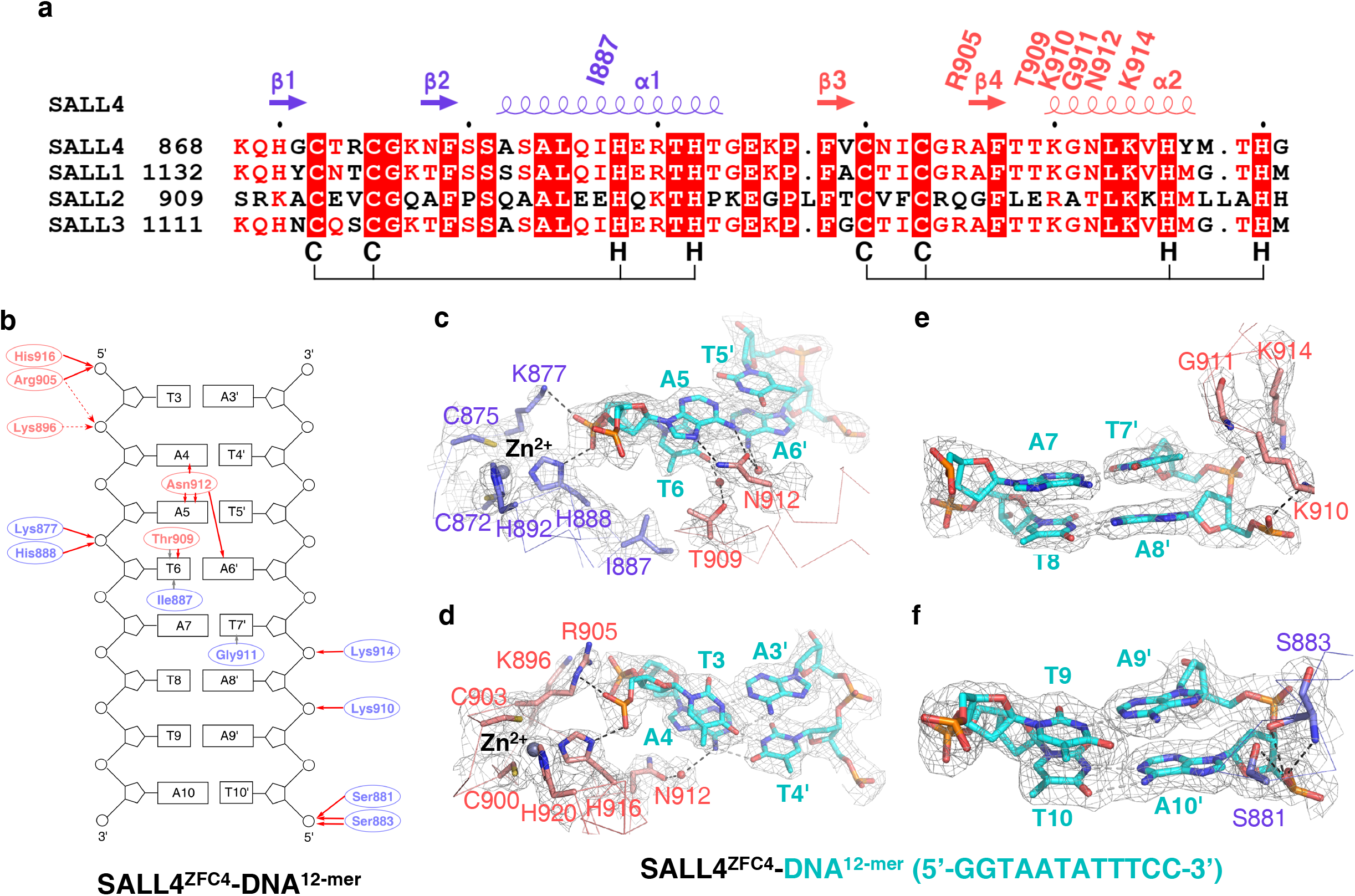
SALL4^ZFC4^ selectively recognizes AT-rich dsDNA. **a** Sequence alignment of human SALL family members, including SALL4 (NP_001304960.1), SALL1 (NP_001121364.1), SALL2 (NP_001278375.1), and SALL3 (NP_741996.2). The secondary structure and DNA binding residues are indicated at the top of sequences, while Zn^2+^ binding residues are indicated at the bottom of sequences. **b** Schematic of the detailed interactions between SALL4^ZFC4^ and DNA. Residues in ZFC4N and ZFC4C are colored in purple and salmon, respectively. Intermolecular hydrogen bonding and hydrophobic interactions are shown in red and grey arrows, respectively. **c-f** Detailed interactions between SALL4^ZFC4^ and **(c)** central ApT (A5-T5’/T6-A6’), **(d)** T3-A3’/A4-T4’, **(e)** A7-T7’/T8-A8’, **(f)** T9-A9’/A10-T10’. Nucleotides are shown in cyan sticks, while ZFC4N and ZFC4C residues involved in DNA binding are shown in purple and salmon, respectively.

SALL4^ZFC4^ wraps around the 12-mer DNA and interacts with it via the positively charged surface (**Fig. 1e-f**). ZFC4N and ZFC4C each belongs to Cys2-His2 (C2H2) finger motif that adopts a canonical β-β-α architecture (**Fig. 2a**). Although the eight Zn^2+^-coordinating residues are absolute conserved in SALL1-4, whereas the spacing between the last two histidines is not conserved in SALL2 (**Fig. 2a**), further suggesting that ZFC4 is not conserved in SALL2 and the C-terminal ZFC of SALL2 probably possesses distinct DNA binding property.

### Structural basis for ^4^AATA^7^-specific recognition by SALL4^ZFC4^

In the structure, the central ^4^AATA^7^ tetranucleotide is recognized by SALL4^ZFC4^ via base-specific hydrogen bonding and Van der Waals interactions (**Fig. 2b**). The SALL4 Asn912 side chain forms two base-specific hydrogen bonds with A5, and another two water-mediated hydrogen bonds with A4 and A6’, respectively (**Fig. 2b-d**). Above Asn-mediated base specific hydrogen bonding interactions were similar to those between Arg and Guanosine observed in the DNA-bound CXXC domain structures ^9, 21^. The side chains of Ile887 and Thr909 make van der Waals interactions with the methyl group of T6, which also forms one water-mediated hydrogen bond with the Thr909 side chain (**Fig. 2b-c**). Furthermore, Gly911 makes Van der Waals interaction with methyl group of T7’, allowing A7 to be favored in the complementary strand (**Fig. 2b and 2e**). Collectively, above hydrogen bonding and van der Waals interactions render SALL4^ZFC4^ the ability to recognize the AATA motif within the 12-mer DNA.

In addition to the base-specific interactions, the SALL4^ZFC4^-DNA complex is further stabilized by extensive electrostatic interactions between DNA backbone and the basic residues of SALL4. The Arg905 and His916 side chains form hydrogen bonds with T3 (**Fig. 2b, 2d**); the side chains of Lys896 and Arg905 make electrostatic interactions with A4 (**Fig. 2b, 2d**); the Lys877 and His888 side chains form two hydrogen bonds with T6 (**Fig. 2b-c**); the side chains of Lys910 and Lys914 form are hydrogen bonded to the backbones of A8’ and T7’ (**Fig. 2b, 2e**), respectively; Ser881 and Ser883 form several hydrogen bonds with the T10’ backbone (**Fig. 2b, 2f**).

Next we applied structure-guided mutagenesis to evaluate the roles of the SALL4 interfacial residues. While N912D abolished the binding, N912A reduced the DNA binding by > 55-fold (KDs: >400 μM vs. 6.9 μM). In contrast, N912Q binds to the DNA with affinity comparable to the wild type (KDs: 8.0 μM vs. 6.9 μM), underscoring the critical role of base-specific hydrogen bonds between Asn912 and A5. The double mutation I887A/T909A weakened the DNA binding affinity by > 10-fold (KDs: 75 μM vs. 6.9 μM), indicating the importance of the Van der Waals interactions between Ile887, Thr909, and Thymine (T6); the triple mutant R905A/K910A/K914A disrupted the DNA binding, indicating the essential role of electrostatic interactions between basic residues and DNA backbone (**Supplementary Table S2**). Collectively, mutagenesis and ITC binding experiments further pinpointed the protein-DNA interface.

### SALL4^ZFC4^ disfavors T or G upstream of ApT

Given that A5 and T6 are engaged in most of base specific interactions, we sought to replace nucleotides flanking A5 to see how it could impact on its DNA binding. All nucleotide replacements were based on the 12-mer dsDNA (5’-^1^GGTAATATTTCC^12^-3’). ITC binding assay demonstrated that T3C/A3’G and A4C/T4’G only slightly weakened the binding to 12-mer AT-rich DNA (KDs: 9.0-11 μM vs. 6.9 μM), whereas A4T/T4’A and A4G/T4’C decreased the SALL4 binding affinity by ∼2.7-5 fold (KDs: 19-36 μM vs. 6.9 μM).

To understand why SALL4 favors AAT and CAT, but not TAT or GAT, we modelled the structures of SALL4 bound with ^4^TAT^7, 4^GAT^7^, and ^4^CAT^7^, respectively (**Fig. 3**). Structural analysis indicates that when A4 is replaced by a Thymine, the distance between the methyl moiety of T4 and Cβ of Asn912 is 3.2 Å, which likely results in the repulsion of the Asn912 side chain and the impaired hydrogen bonds between Asn912 and A5 (**Fig. 3a-b**). In addition, A4T disrupts the water-mediated hydrogen bond between Asn912 and A4 (**Fig. 3b**). In the modelled structure of SALL4 bound with ^4^GAT^7^, N7 and O6 of G4, and carboxyl oxygen of Asn912, are all hydrogen bond acceptors, which disrupts the water-mediated hydrogen bond observed between Asn912 and A4 (**Fig. 3C**). In contrast, A4C did not affect the hydrogen bond between Asn912 and A4, and also maintains the water-mediated hydrogen bond (**Fig. 3D**). However, if C4 is methylated to mC4, mC4 would weaken the Asn912-A5 hydrogen bond as T4 does. Thus, our structural analysis and binding data further reveal that SALL4 prefers an ^4^AATA^7^ or a ^4^CATA^7^motif within the 12-mer dsDNA.

**Figure 3.**
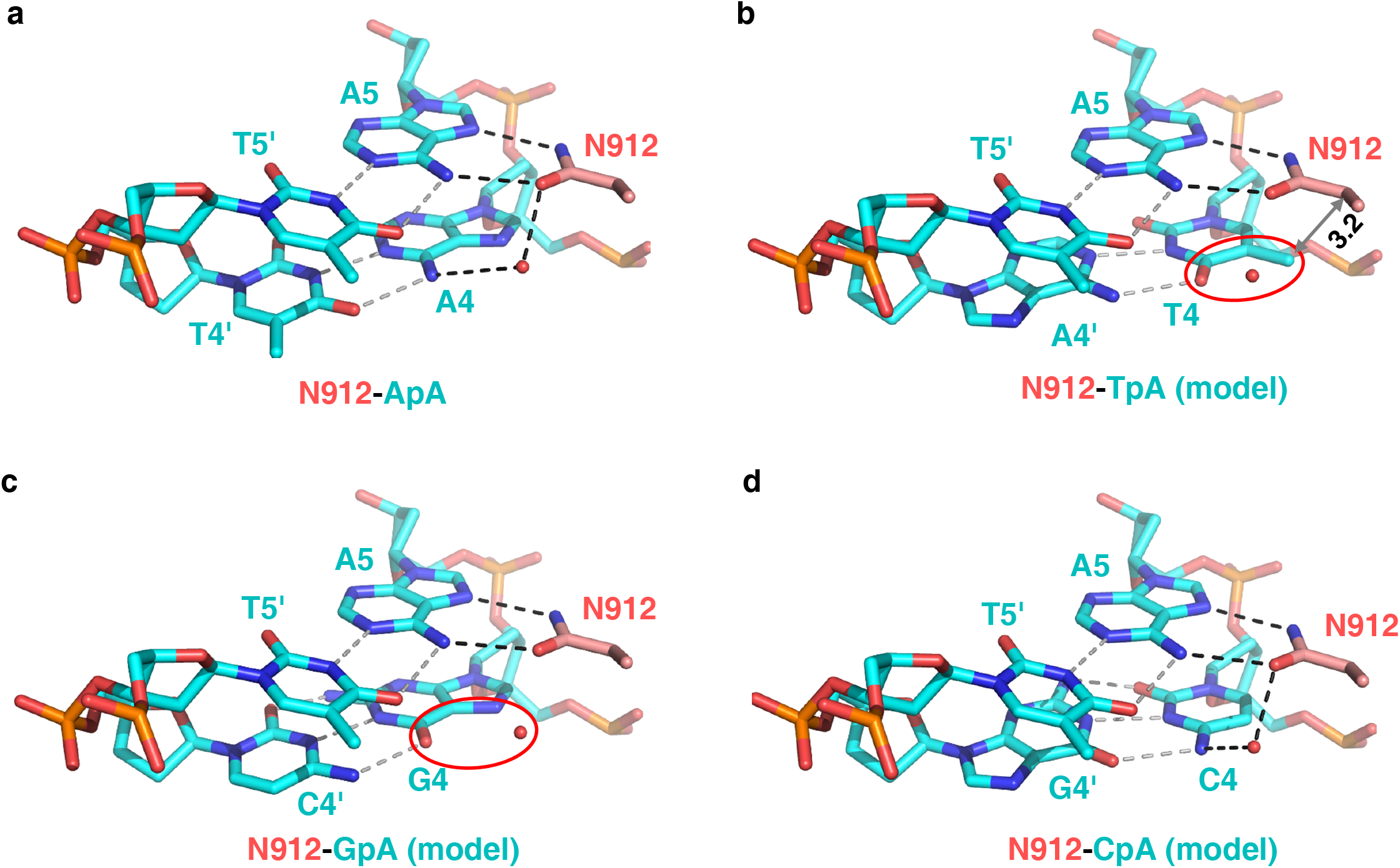
SALL4^ZFC4^ Asn912 disfavored TpA and GpA dinucleotides. **a** In the DNA-bound SALL4^ZFC4^ structure, A5 is recognized by Ans912, which further stacks with upstream A4. **b** In the modeled structure, the A4 substituted by T4 leads to potential steric clash between Asn912 side chain and the methyl group of T4. **c** The A4 substituted by G4 disrupts the water-mediated hydrogen bond. **d** The A4 substituted by C4 maintains the Asn912-mediated base specific interactions.

To study whether the AT-rich recognition by SALL4^ZFC4^ also applies for dsDNAs of different lengths, we determined the 2.5 Å structure of SALL4^ZFC4^ with a 16-mer dsDNA containing two ATA motifs (**Supplementary Table S2**). In the structure, the two SALL4^ZFC4^ molecules recognize ^8^AATA^11^ and ^5^TATA^8^ within the 16-mer dsDNA, respectively, to form the complex in a 2:1 molar ratio (protein: DNA) (**Supplementary Fig. S1a-e**). The sequence-specific recognition mode is the same as observed in the 12-mer DNA complex. Consistently, the ITC binding assay also demonstrates that SALL4^ZFC4^ binds to the 16-mer dsDNA with two KDs in a range of 13-16 μM. While N912Q mutant binds to the 16-mer DNA with KDs similar to those of the wild type (KDs: 16-19 μM vs. 13-16 μM), R905A/K910A/K914A mutant displays no binding towards the 16-mer dsDNA (**Supplementary Table S1**). Thus, we conclude that SALL4^ZFC4^ specifically recognizes the AT-rich motif within dsDNAs, regardless of the DNA length.

### Structure of SALL3^ZFC4^ with the 12-mer AT-rich dsDNA

To understand whether above DNA recognition mode also applies for other SALL members, we determined the crystal structure of SALL3^ZFC4^ with the same 12-mer AT-rich dsDNA in a 2.50 Å resolution (**Supplementary Table S2**). There is only one SALL3^ZFC4^-DNA complex in an asymmetric unit. Similar to that of SALL4 ^ZFC4^ with 12-mer dsDNA, SALL3^ZFC4^ binds to the central major groove of the 12-mer DNA via its extensive positive charged surface (**Fig. 4a**). The DNA-bound SALL3^ZFC4^ structure is superimposed well with the two SALL4 complexes (**Fig. 4b**), suggesting the conserved architecture of SALL complexes.

**Figure 4.**
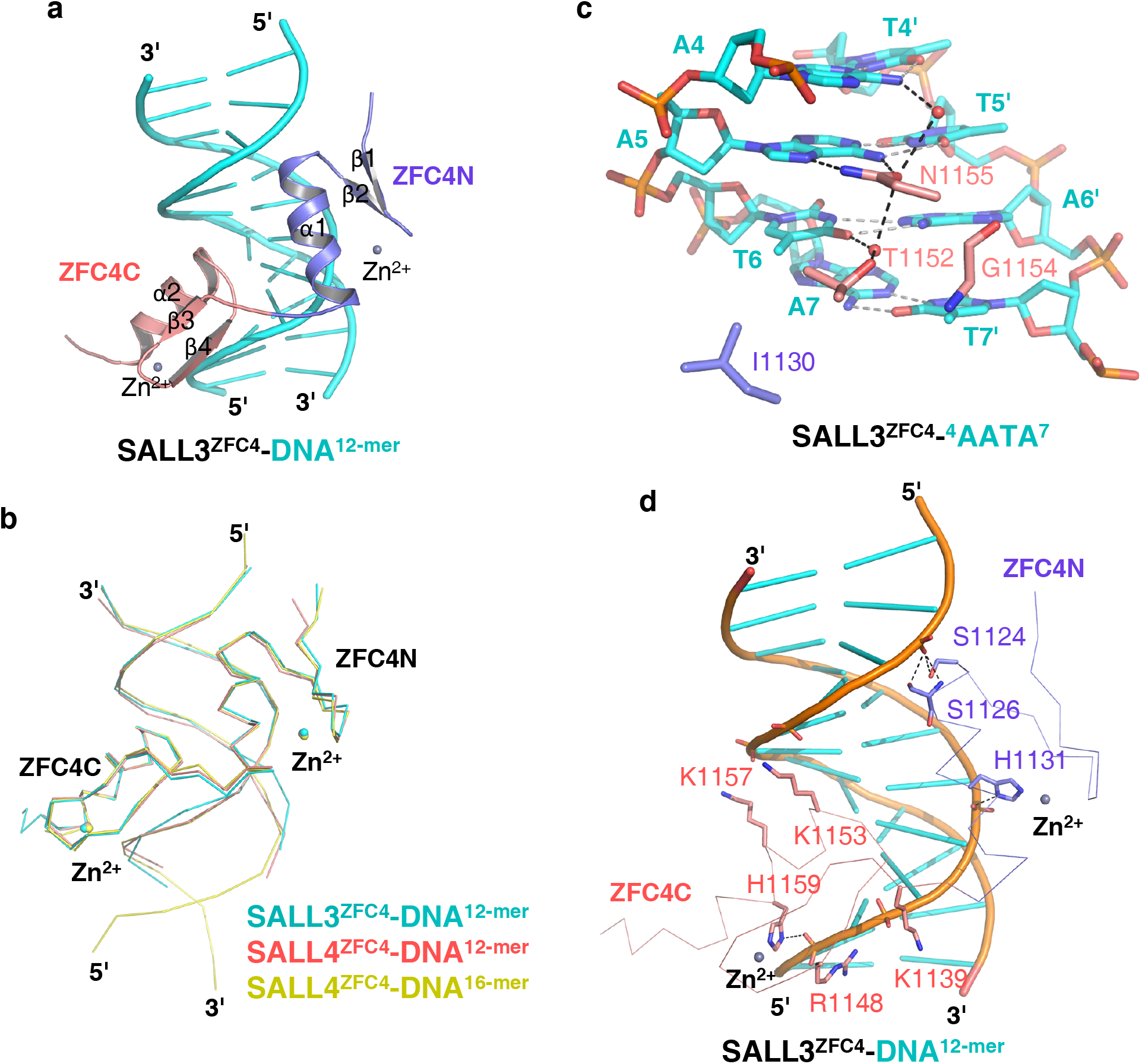
SALL3^ZFC4^ specifically recognizes AT-rich dsDNA. **a** Structure of SALL3^ZFC4^ bound with the 12-mer dsDNA (5’-GGTAATATTTCC-3’), which is colored the same as in Fig. 1E. **b** Superposition of the structures of SALL3^ZFC4^ with the 12-mer dsDNA (cyan ribbon), SALL4^ZFC4^ with the 12-mer dsDNA (red ribbon), and SALL4^ZFC4^ with the 16-mer dsDNA (yellow ribbon). **c** Base specific interactions between SALL3^ZFC4^ and central ^4^AAT^6^, which are colored the same as in Fig. 1e. **d** Interactions between SALL3^ZFC4^ and DNA backbone. Protein and DNA are shown in ribbon and cartoon, respectively. SALL3 residues are colored the same as in Fig. 1e.

Extensive hydrogen bonding and Van der Waals interactions were found between SALL3^ZFC4^ and DNA. SALL3 Asn1155, the counterpart of SALL4 Asn912, forms two base-specific hydrogen bonds with A5; Ile1130 and Thr1152 of SALL3, the counterparts of SALL4 Ile887 and Thr909, respectively, make Van der Waals interaction with T6, which forms a water-mediated hydrogen bond with Thr1152; Gly1154 makes additional Van der Waals interaction with T7’ (**Fig. 4c**). Overall, ^4^AATA^7^ recognition by SALL3^ZFC4^ is the same as that by SALL4^ZFC4^. In addition, the electrostatic interactions between DNA backbone phosphates and SALL3 residues, including Ser1124, Ser1126, His1131, Lys1139, Arg1148, Lys1153, Lys1157, and His1159, are also conserved in the SALL4 complex (**Fig. 4d**). Since the DNA binding residues of SALL3 are conserved in SALL1 but not SALL2, we reason that the AT-rich DNA recognition is conserved in ZFC4 of SALL1, SALL3, and SALL4.

### SALL4 ^ZFC1^ recognizes AT-rich DNAs

Sequence alignment of SALL4 ZFC1 and ZFC4 shows that all SALL4^ZFC4^ residues involved in the recognition of AT-rich DNA are conserved in SALL4^ZFC1^ except Ala882, which is replaced by an Asp (Asp394) in SALL4^ZFC1^ (**Supplementary Figure S2A**). Then we examined the DNA binding of SALL4^ZFC1^ (aa 378-453) by ITC. Binding data show that SALL4^ZFC1^ binds to the 12-mer AT-rich DNA with a KD of 24 μM, and binds to the 16-mer AT-rich DNA with KDs in a range of 17-21 μM (**Supplementary Table S1**), weaker than that for SALL4^ZFC4^.

We further solved the structure of SALL4^ZFC1^ with 16-mer AT-rich DNAs in a 2.72 Å resolution (**Supplementary Table S2**). There are three dsDNAs and six SALL4^ZFC1^ molecules in one asymmetric unit, with one dsDNA bound with two SALL4^ZFC1^ molecules (**Fig. 5a**). In the structure, all six SALL4^ZFC1^ recognizes A-T base pair (A9-T9’ or A7’-T7) via its Asn424 (**Fig. 5b-g**). Asn424 of molecules A, D, and E recognizes A9 in the context of ApA (**Fig. 5b, 5e, and 5f**), whereas Asn424 of molecules B, C, and F interacts with A7’ in the context of TpA (**Fig. 5c, 5d, and 5g**). The lengths of hydrogen bonds between Asn424 and A7’ are in a range of 3.1-3.6 Å, longer than those observed between Asn424 and A9 (2.7-3.1Å) (**Fig. 5b-g**), suggesting that the hydrogen bonds are weakened by the upstream Thymine (T8’).

**Figure 5.**
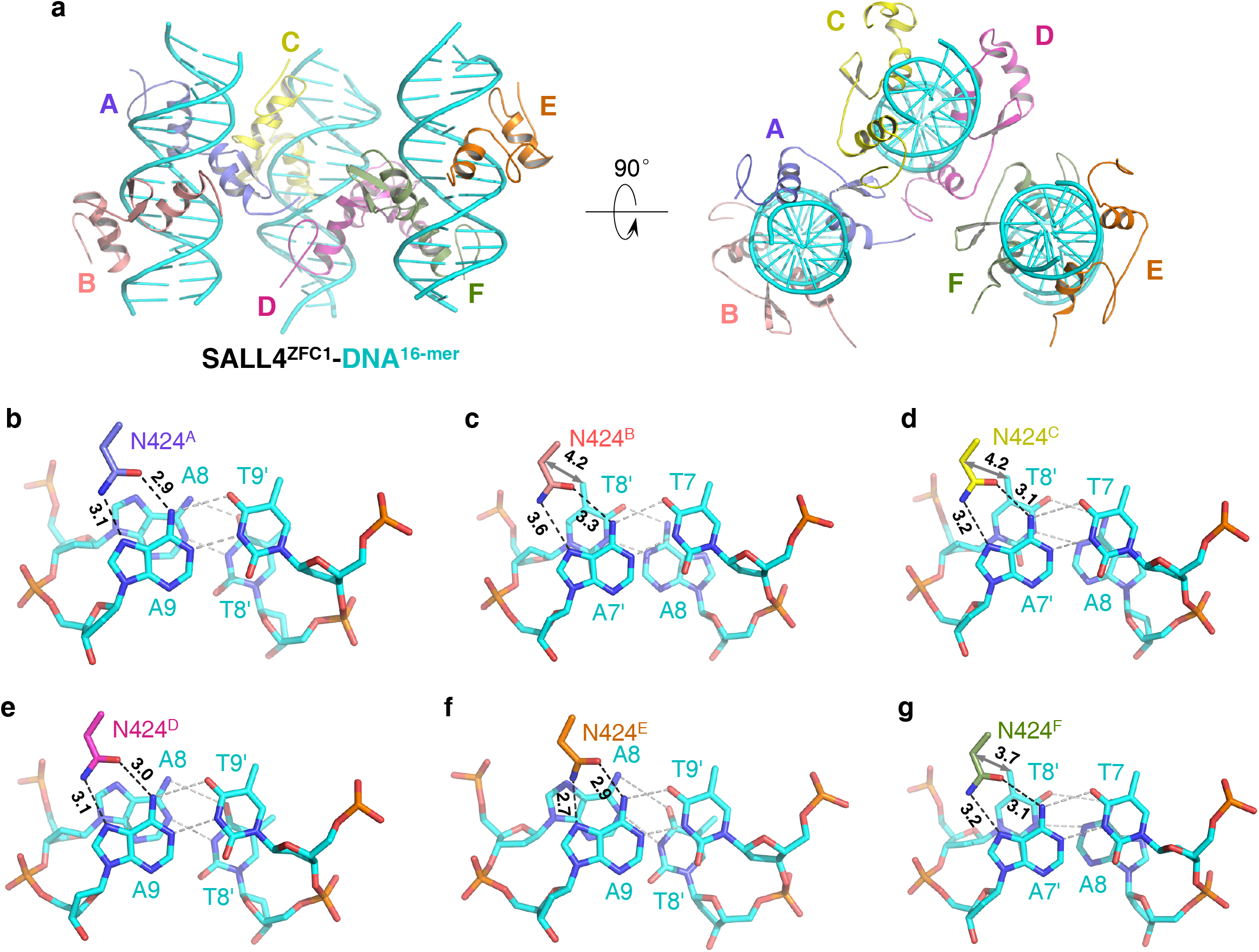
SALL4^ZFC1^ selectively binds to AT-rich dsDNA. **a** Structure of SALL4^ZFC1^ bound with the 16-mer dsDNA (5’-GGAATATAATATTTCC-3’). Three dsDNAs are shown in cyan cartoon, while six SALL4^ZFC1^ molecules are shown in cartoon with different colors. **b-g**, all six SALL4^ZFC1^ molecules recognizes central adenosine via Asn424. In b, e, and f, Asn424 recognizes A9 in the context of ApA; In c, d, and g, Asn424 recognizes A7’ in the context of TpA. Asn424 is shown in sticks and all DNAs are shown in cyan sticks.

Next we superimposed the structure of SALL4^ZFC1^ with the 16-mer dsDNA with that of SALL4^ZFC4^ with the same DNA, and found that Ala882 of ZFC4 is spatially proximal to the phosphate group of A14’ due to the hydrogen bond between the DNA backbone and the main chain amide of Ser883 (**Supplementary Fig. S2A**). In contrast, SALL4^ZFC1^ Asp394, the counterpart of SALL4^ZFC4^ Ala882, leads to charge repulsion with the DNA backbone phosphate, which would impair the hydrogen bond between A14’ and SALL4^ZFC1^ Ser395, the counterpart of SALL4^ZFC4^ Ser883 (**Supplementary Fig. S2B**). Consistent with the structural analysis, we found that A882D of SALL4^ZFC4^ reduced the DNA binding affinity by > 6.5-fold (KDs: 47 μM vs. 6.9 μM) (**Supplementary Fig. S2C**). Collectively, our structural data, complemented by mutagenesis and binding experiments, reveals that SALL4^ZFC1^ also specifically recognizes AT-rich DNA, albeit with weaker affinity.

### Targeting of SALL4 at AT-rich sites inhibits aberrant expression of differentiation prompting genes

In mouse ESCs, binding of SALL4 to AT-rich putative enhancer sequences prevents expression of differentiation promoting genes ^14^. To test whether loss of DNA binding in the SALL4^ZFC4^ Asn912 mutation has a biological significance, we generated *Sall4*^*-/-*^ mouse ESCs from *Sall4*^*-/flox*^ ESCs by infecting adenovirus-EGFP-Cre, which does not integrate into the genome. Then, we infected *Sall4*^*-/-*^ ESCs with lentivirus carrying either wild-type (WT) mouse *Sall4* or mouse *Sall4* N922D mutant (human SALL4 Asn912 corresponds to mouse SALL4 Asn922). First, we examined *Sall4* expression levels by qRT-PCR. Expression of transgene WT *Sall4* and *Sall4* N922D is approximately 1.8 and 0.9 fold, respectively, compared to *Sall4* expression in control ESCs (**Fig. 6a**). Then, we examined expression of several neural differentiation genes to which SALL4 is enriched (**Fig. 6c, g**).

**Figure 6.**
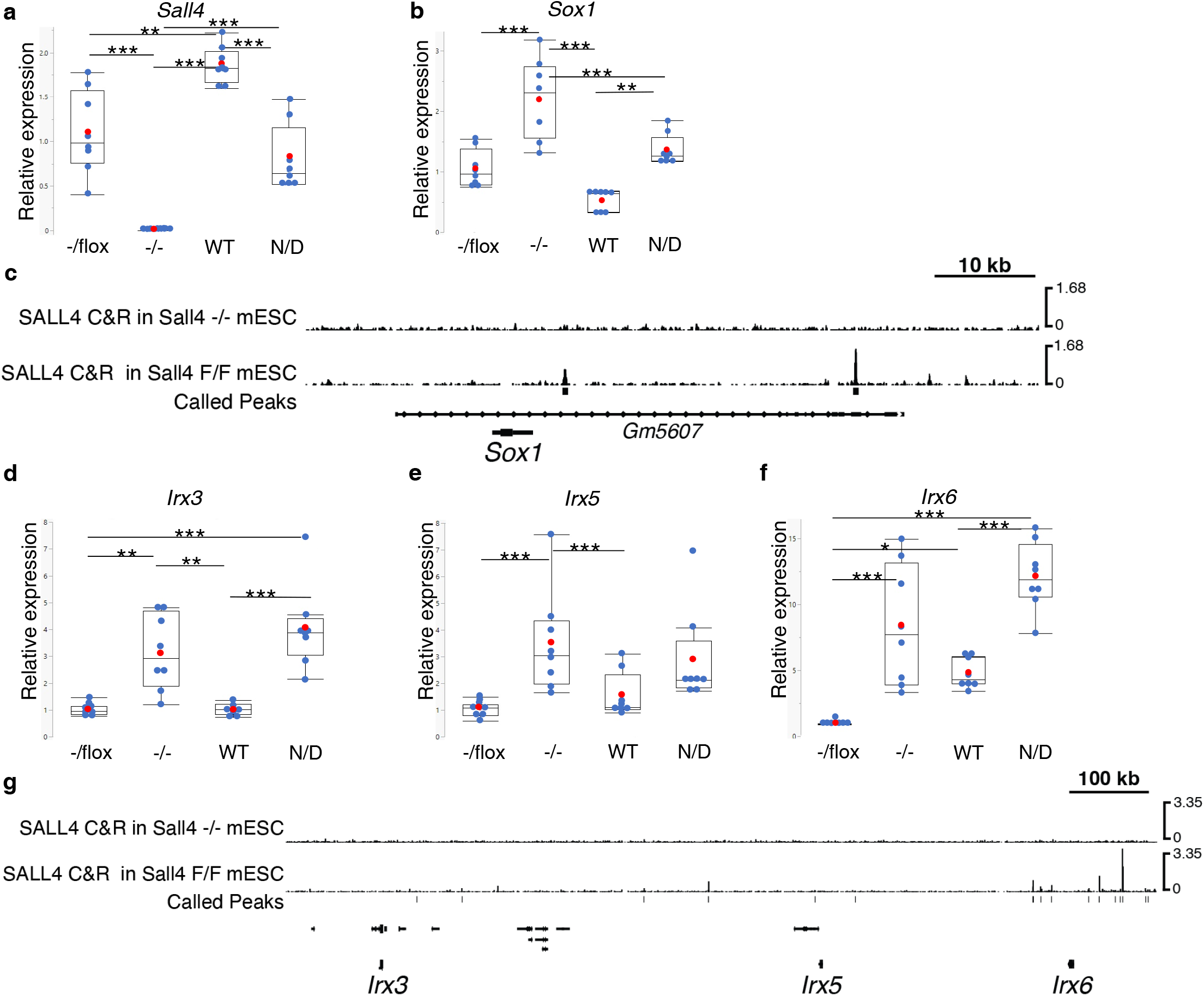
*mSall4* N922D mutant partially rescues aberrant gene expression in *Sall4*^-/-^ mouse ESCs. (**a, b, d-f**) Box plots showing relative expression levels of *mSall4* (**a**), *Sox1* (**b**), *Irx3* (**d**), *Irx5* (**e**) and *Irx6* (**f**) in *Sall4*^-/flox^ cells, *Sall4*^-/-^ cells, *Sall4*^-/-^ cells with WT *mSall4* expression (WT) and *Sall4*^-/-^ cells with *Sall4* N922D (ND) mutant expression. *: p<0.05, **: p<0.01, ***: p<0.001 by One-Way ANOVA with post-hoc Tukey HSD test. Each replicate is shown as a blue dot, and the average value is shown as a red dot. **(c, g)** SALL4 CUT&RUN tracks of the *Sox1* and *Irx3*-*Irx5*-*Irx6* regions in *Sall4*^-/-^ and *Sall4*^flox/flox^ (F/F) mouse ESCs. In g, genes other than *Irx3*-*Irx5*-*Irx6* are not labeled for the simplicity.

By CUT&RUN experiments, we observed specific enrichment of SALL4 near the *Sox1* gene (**Fig. 6c**). As previously shown ^20^, *Sox1* expression is elevated in *Sall4*^-/-^ ESCs, compared to the control (**Fig. 6b**). Both WT *Sall4* and *Sall4* N922D, introduced into *Sall4*^*-/-*^ ESCs, prevented aberrant expression of *Sox1* (**Fig. 6b**). SALL4 was also enriched in the region where *Irx3, Irx5* and *Irx6* are closely located on chromosome 8 (**Fig. 6g**). Expression of *Irx3* was elevated in *Sall4*^-/-^ ESCs. The WT *Sall4* transgene repressed aberrant expression of *Irx3*, but *Sall4* N922D failed to repress *Irx3* expression (**Fig. 6d**). Similarly, WT, not N922D *Sall4* repressed aberrant expression of *Irx5* (**Fig. 6e**), and a similar trend was observed for *Irx6* (**Fig. 6f**). In agreement with a previous study^14^, AT-rich sequences are enriched in the called peaks in the CUT&RUN experiments. Specifically, both peaks near the *Sox1* gene and 15 out of 17 peaks at the *Irx3-Irx5-Irx6* region contain the AATA sequence, which was found in 55,995 of the 64,249 (87%) of the genome-wide SALL4 binding peaks. Inhibition of aberrant expression of *Sox1* and *Irx* genes is consistent with the notion that ZFC4-dependent DNA binding of SALL4 contributes to repression of these gene expression. In addition, repression of *Sox1* by *Sall4* N992D might be associated with the binding of SALL4 N922D to the AT-rich region via its ZFC1 domain, consistent with our structural and biochemical data.

## Discussion

More than 700 Zinc fingers proteins in human genome belong to the C2H2-type, and ∼400 of them were annotated as TFs ^22, 23^. Uncovering the preferred motif of zinc finger TFs is important for understanding their roles in orchestrating spatiotemporal gene transcription. SALL family members are a subfamily of zinc finger proteins playing important roles in cell development and differentiation. Dysfunctional SALL proteins are associated with different types of cancers. In this study, we uncovered the conserved AT-rich DNA recognition mode by SALL family members through presenting several structures of SALL proteins with respective DNA ligands. SALL proteins utilizes a conserved Asn, such as Asn912 of SALL4 or Asn1155 of SALL3, to recognize the Adenosine of A-T base pair, while hydrophobic residues of SALL proteins interact with the methyl moiety of the downstream Thymine. In addition, the adenosine binding Asn also favors an Adenosine at upstream position. Overall these base-specific interactions confer SALL proteins the ability to interpret AT-rich DNAs.

Given that SALL4^ZFC1^ binds to AT-rich DNAs weaker than SALL4^ZFC4^, it might be insufficient to maintain the occupancy of SALL4 after the deletion of ZFC4, consistent with previous report that mutation or deletion of SALL4^ZFC4^ impairs the targeting of SALL4 at genome sites^14^.

### Comparison of SALL4-DNA structure with those of other AT-rich DNA complexes

It has been reported that transcriptional repressor MogR specifically recognizes AT-rich DNAs^24^, which prompts us to compare it with the SALL4^ZFC4^ complex. The DNA recognition mode in two complexes are quite different (**Supplementary Fig. S3**). MogR specifically recognizes the AT-rich motif, but in a manner distinct from that between SALL4 and DNA ligands (**Supplementary Fig. S3**). Asn118 of MogR forms two hydrogen bonds with A6’ and A5’ (**Supplementary Fig. S3B**), respectively, different from the Asn-Adenosine pair observed in SALL4 complex (**Supplementary Fig. S3A**); Ser114 and Gln117 forms water-mediated hydrogen bonds with A5’ and T4, respectively; Methyl groups of T4 and T3 make hydrophobic interactions with Tyr121 and Val94, respectively (**Supplementary Fig. S3B**) ^24^. Therefore, above interactions renders MogR the ability to recognize ^3^TTTT^6^, which is different from those observed in SALL4 complexes.

### Disease associated mutations

In contrast with many deletions in SALL4 that were known to result in Okihiro Syndrome. Only very few single mutations within ZFC4, including H888R, is reported to be associated with Okihiro Syndrome. Based on the Catalogue of Somatic Mutations in Cancer (COSMIC) database (https://cancer.sanger.ac.uk/cosmic), identified single mutations in SALL4^ZFC1^ and SALL4^ZFC4^ that likely have impact on protein stability or include S396F and R431Q of SALL4^ZFC1^, and H888Q, R905Q, K914N, and H916Y of SALL4^ZFC4^. S396F disrupts the intramolecular hydrogen bond, and might destabilize the protein, while R431Q of SALL4^ZFC1^ weakens the interaction with DNA backbone; R905Q and K914N weaken the binding of SALL4^ZFC4^ with DNA backbone, whereas H888Q and H916Y not only disrupt the binding to Zn^2+^, but also abolish the hydrogen bond with DNA backbone. These disease associated mutations indicate that impaired DNA binding affinities of SALL4^ZFC1^ and SALL4^ZFC4^ might be associated with human cancers.

The N-terminal 12 amino acid stretch of SALL4 interacts with the nucleosome remodeling deacetylase (NuRD) complex that creates repressive chromatin structure in ESCs^25, 26^. Our structural study illustrates a module-specific roles of SALL4 in target sequence recognition by ZFC and recruiting NuRD. In this way our study not only uncovers the conserved DNA recognition mode by SALL family members, but also provides insights into a better understanding how SALL4 mutations result in human cancers via altering the expression profile of key regulators such as *Sox1*.

In sum, our study not only provides mechanistic insight into the AT-rich DNA recognition by SALL4^ZFC4^ and SALL4^ZFC1^, but also uncovers that the binding of SALL4 at specific AT-rich genomic DNA regions influences cell differentiation and cell fate *in vivo*.

## METHODS

### Cloning, protein expression and purification

Gene encoding SALL4^ZFC4^ (residues 856-930) was amplified by PCR from a complementary DNA library; Genes encoding human SALL4^ZFC1^ (aa 378-453) and SALL3^ZFC4^ (aa 1102-1166) were synthesized by genscript (Nanjing); Gene encoding human SALL4^548-1029^, which spans ZFC2 and ZFC4, was synthesized by Sangon Biotech (Shanghai). All of them were cloned into pET28-MHL vector, and the cloned plasmid was transformed into *E. coli* BL21 (DE3). Cells were grown in LB medium at 37 °C until the OD600 reached ∼0.8. The recombinant protein was overexpressed at 16 °C for 18h after induction by 0.2 mM (final concentration) β-D-1-thiogalactopyranoside (IPTG) and 40 μM ZnCl2. Cells were harvested by centrifuge at 3600 × g, 4 °C for 15 min and pellets were resuspended in a buffer containing 20 mM Tris-HCl, pH7.5, and 400 mM NaCl. Lysates were centrifuged at 10000 × g, 4 °C for 30 min and supernatants were collected.

Recombinant SALL4^ZFC4^ was purified by Ni-NTA column (GE healthcare), and eluted by 20 mM Tris-HCl, pH7.5, 400 mM NaCl and 500 mM imidazole. N-terminal polyhistidine tags (His-tags) of the recombinant protein was cleaved by Tobacco etch virus (TEV) protease and dialyzed overnight with the buffer containing 20 mM Tris-HCl, pH7.5, and 150 mM NaCl. SALL4^ZFC4^ was further purified by Superdex 75 gel filtration (GE Healthcare) and HitrapTM S HP column (GE healthcare). The purified protein was concentrated to 8 mg/ml in the buffer containing 20 mM Tris-HCl, pH7.5, and 150 mM NaCl, and was stored at −80 °C before further use.

Expression and purification of SALL4^ZFC1^ and SALL3^ZFC4^ were performed in the same way as for SALL4^ZFC4^. Site-specific mutations were carried out by using two reverse and complement primers containing the mutation codon. Primer sequences used for cloning mutants are listed in **S**upplementary Table S3. All mutants were purified in the same way as for wild type proteins.

### Isothermal titration calorimetry (ITC)

ITC experiments were performed on a MicroCal PEAQ-ITC calorimeter (Malvern Panalytical) at 25 °C by titrating 2 μl of protein (2 mM) into cell containing 40 μM double strand DNA, with a spacing time of 120 s and a reference power of 5 μCal s^−1^. Control experiments were performed by titrating protein (1mM) into buffer only, and were subtracted during analysis. Binding isotherms were plotted, analyzed and fitted by MicroCal PEAQ-ITC Analysis software (Malvern Panalytical). The dissociation constants (KDs) were determined from a minimum of two experiments (mean ± SD). dsDNAs used for ITC are listed in Supplementary Table S4.

The mutant protein SALL4^ZFC4^ N912Q mutant is less stable under the condition of 20 mM Tris-HCl, 150 mM NaCl, pH7.5, and its ITC experiments were carried out by titrating 2 μl of double strand DNA (0.7 mM) into cell containing 40 μM proteins. Representative ITC binding curves are shown in Supplementary Figure S5.

### Crystallization, data collection and structure determination

All crystals were grown using the sitting drop vapor diffusion method at 18 °C. Before crystallization, protein is mixed with dsDNA ligand at a ratio of 1:1. Crystal of SALL4^ZFC4^ in complex with the 12-mer dsDNA (5’-GGTAATATTTCC-3’) was obtained by mixing 1.0 μl of complex with 1.0 μl of well solution containing 0.1 M BIS-TRIS, pH6.5, and 25% (w/v) polyethylene glycol 3350 (PEG 3350). Crystal of SALL4^ZFC4^ in complex with the 16-mer dsDNA (5’-GGAATATAATATTTCC-3’) was obtained by mixing 1.0 μl of complex with 1.0 μl of well solution containing 0.1 M BIS-TRIS, pH 5.5, 0.2 M Sodium chloride, and 25% PEG 3350. Crystal of SALL4^ZFC3^ in complex with the 12-mer dsDNA (5’-GGTAATATTTCC-3’) was obtained by mixing 1.0 μl of complex with 1.0 μl of well solution containing 0.1 M MES monohydrate, pH 6.5, 0.2 M Ammonium sulfate, and 30% w/v polyethylene glycol monomethyl ether 5000. Crystal of SALL4^ZFC1^ in complex with the 16-mer dsDNA (5’-GGAATATAATATTTCC-3’) was obtained by mixing 1.0 μl of complex with 1.0 μl of well solution containing 0.1M HEPES, pH6.5, 10% PEG 6000, and 5% (v/v) 2-Methyl-2,4-pentanediol (MPD). Before flash-freezing crystals in liquid nitrogen, crystals were soaked in a cryoprotectant consisting of 85% reservoir solution plus 15% glycerol.

The diffraction data were collected on beam line BL17B and BL18U1 at Shanghai Synchrotron Facility (SSRF), and processed with HKL2000/3000 ^27, 28^ or XDS ^29^. For the complex structure of SALL4^ZFC4^ with 12-mer DNA, the initial model of SALL4^ZFC4^ was solved by CRANK2 ^30^ using the zinc single-wavelength anomalous dispersion phasing. The DNA was built manually by COOT ^31^, and the complex model was further refined by Phenix^32^. The other complexes were solved by molecular replacement using Phaser^33^ with previously solved SALL4^ZFC4^ complex as the search model. Then the models were built and refined manually by COOT^31^, and were further refined by Phenix ^32^. The statistic details about data collection and structure refinement were summarized in Supplementary Table S2.

## Mouse ESC culture

*Sall4*^-/flox^ mouse ESCs were previously described ^34^. Cell are maintained in the 2i media ^35^. To generate *Sall4*^-/-^ cells, *Sall4*^-/flox^ ESCs were suspended by trypsinization and neutralization, then, infected with adenovirus EGFP-Cre ^36^. Independent clones were isolated, expanded, and the *Sall4*^*-/-*^ genotype was confirmed by genomic PCR as previously described ^34^.

Wild type mouse *Sall4* was cloned in the pLV-EF1a-IRES-Puro vector ^37^. The *Sall4* N922D mutant was generated by site directed mutagenesis using Q5 High-Fidelity DNA Polymerase (New England Biolabs) and In-Fusion Snap Assembly (Takara Bio USA) following the manufacturer’s instructions. Lentiviruses were produced according to a standard procedure ^38^, and were concentrated using Lenti-X Concentrator (Takara Bio USA). Approximately, 1× 10^5^ *Sall4*^*-/-*^ mESCs were infected with lentivirus carrying WT or mutant *Sall4* and selected by 2μg/mL puromycin. Selected cells were expanded and used for experiments.

For qRT-PCR, total RNA was isolated using the Direct-zol RNA MicroPrep kit (Zymo Research) and cDNA was synthesized using iScript cDNA synthesis kit (BioRad) according to the manufacturers’ instructions. qPCR was performed using SYBR green master mix (ThermoFisher) and primers in Supplemental Table 5.

## CUT&RUN experiments

CUT & RUN ^39^ was performed essentially as described in the online protocol (dx.doi.org/10.17504/protocols.io.zcpf2vn) using *Sall4*^*flox/flox*^ or *Sall4*^*-/-*^ mouse ESCs (10^5^ cells per reaction) cultured in the 2i + LIF media ^20^. Anti-SALL4 antibody (SC-101147 (EE-30)) or normal rabbit IgG (SC-2025) were each used at a 1:300 dilution. EDTA was excluded from all buffers prior to MNase inactivation to avoid Zn+ chelation. Cell permeabilization and all subsequent steps were performed using buffers containing 0.02% digitonin. Recovered DNA fragments were end-repaired, A-tailed and ligated with xGen adapters (10005974, Integrated DNA Technologies) using the Kapa Hyper Prep Kit (07962312001, Roche) and barcoded during amplification using Kapa HotStart Readymix (7958927001, Roche). Libraries were sequenced using a 2 × 150 paired-end configuration on a HiSeq 4000 (Genewiz). Reads were trimmed using TrimGalore (0.6.0) and CutAdapt (1.18) and read quality was assessed with Fastqc (0.11.8). Trimmed reads were mapped with BWA MEM (0.7.17) using the mouse genome (GRCm38) as reference. Peaks were identified using MACS (2.1.1.20160309) using the --call-summits -g mm parameters. Peak lists from each replicate were merged using R (4.1.2) to find high confidence peaks present in both replicates. Raw and processed data files are available in the Gene Expression Omnibus using accession GSE203303.

## Data availability

The coordinates and structure factors files for the structures of SALL4^ZFC4^ with 12-mer dsDNA, SALL4^ZFC4^ with 16-mer dsDNA, SALL3^ZFC4^ with 12-mer dsDNA, and SALL4^ZFC1^ with 16-mer dsDNA, were deposited into Protein Data Bank under accession codes 7Y3I, 7Y3K, 7Y3L, and 7Y3M, respectively.

## Acknowledgements

We thank the staff from the BL17B/BL18U1/BL19U1/BL19U2/BL01B beamline45 of National Facility for Protein Science in Shanghai at Shanghai Synchrotron Radiation Facility for assistance during data collection. This work is supported by the National Natural Science Foundation of China (grant nos. 22137007 and 92053107). C.X. is supported by “the Fundamental Research Funds for the Central Universities”, the Major/Innovative Program of the Development Foundation of the Hefei Center for Physical Science and Technology (2021HSC-CIP014), and “the Thousand Young Talent program”. Y.K is supported by a grant from National Institutes of Health of USA (R01AR064195) and Grant-in-Aid of Artistry, Research and Scholarship of the University of Minnesota (#378836).

